# Sex-Specific Roles of Hypocretin Receptor Signaling in CRF Neurons on Alcohol Drinking, Anxiety, and BNST Neuronal Excitability

**DOI:** 10.1101/2024.09.07.609774

**Authors:** Yihe Ma, Haniyyah Sardar, Max E. Benabou, Angeline C. Yu, Allison R. Morningstar, R. Nicolas Fajardo, Isaac F. Kandil, Ethan T. Rogers, Anne Vassalli, Julie A. Kauer, William J. Giardino

## Abstract

Alcohol use disorder (AUD) is characterized by compulsive alcohol consumption and negative emotional states during withdrawal, often perpetuating a cycle of addiction through arousal dysfunction. The hypocretin/orexin (Hcrt) neuropeptide system, a key regulator of arousal, has been implicated in these processes, particularly in its interactions with corticotropin-releasing factor (CRF) neurons within the bed nucleus of the stria terminalis (BNST). We investigated the role of Hcrt receptor signaling in CRF neurons in modulating alcohol intake, anxiety behaviors, and BNST excitability, with a focus on sex-specific differences. Using CRF-specific genetic deletion of HcrtR1 and/or HcrtR2 receptors in mice, we found that deletion of HcrtR1 significantly reduced alcohol intake, with sex-specific effects on BNST excitability. CRF-specific HcrtR2 deletion, while not affecting alcohol consumption, decreased baseline anxiety-like behaviors in males relative to females. Moreover, the double deletion of both Hcrt receptors from CRF neurons led to reduced alcohol drinking in males and dampened anxiety behaviors and BNST excitability in both sexes during protracted withdrawal. These findings suggest that Hcrt signaling in CRF neurons plays a critical role in the persistence of excessive alcohol consumption and the development of negative affective states, with distinct contributions from HcrtR1 and HcrtR2. The observed sex-specific differences underscore the need for tailored therapeutic approaches targeting the Hcrt system in the treatment of AUD.

## Introduction

Alcohol use disorder (AUD) is characterized by a compulsion to seek and drink alcohol, an inability to control alcohol consumption, and the predominance of negative emotional states during withdrawal when alcohol is unavailable^1^. Affective disturbances are highly comorbid with AUD and contribute to initial alcohol use, excessive, alcohol consumption, and long-term neural adaptations occurring as a consequence of chronic alcohol exposure^2^. Specifically, AUD can generate maladaptive neuroplasticity leading to arousal dysfunction, such as sleep disruption and emotional dysregulation (namely, heightened anxiety)^3^. In turn, arousal dysfunction exacerbates the key characteristics of AUD, perpetuating the addiction cycle^4^. Growing interest in the study of “hyperkatifeia” (enhanced sensitivity to the negative emotional symptoms of addiction during withdrawal) has led to increased focus on identifying the mechanisms by which emotional centers of the limbic system promote negative affect in AUD^5,6^. Given the complexity of cell types, neuromodulators, and pathways that interact with unique motivational centers in the brain^7–13^, this challenge has required advanced technological and multidisciplinary approaches.

The hypocretin/orexin (Hcrt) neuropeptide system is a key modulator of the brain’s wakefulness/arousal circuitry^14–16^. Through its interactions with other neural systems, Hcrt activity and signaling maintains adequate levels of arousal despite fluctuations in internal and external states^17^. Thus, dysfunction of Hcrt system activity can destabilize arousal regulation, enhancing susceptibility to AUD^7,9,18–22^. Hcrt-expressing neurons are located in the lateral hypothalamic area (LH) and produce prepro-orexin precursors, which are cleaved into two Hcrt neuropeptides (Hcrt1/orexin-A and Hcrt2/orexin-B)^17,23^. Hypocretins act through two G-protein-coupled receptors (HcrtR1 and HcrtR2), which signal through Ca2+/cAMP stimulatory Gq-coupled or Gs-coupled pathways, with HcrtR1 displaying a higher affinity for Hcrt1, while HcrtR2 has similar affinities for both peptides^24^. The two receptor subtypes are differentially expressed throughout the brain, suggesting they serve diverse physiological functions, although both receptors have been identified in areas including the bed nucleus of the stria terminalis (BNST), cortex, nucleus accumbens, paraventricular thalamus, central amygdala, and ventral tegmental area^25,26^. These regions overlap with substrates involved in AUD, and differential HcrtR signaling patterns throughout this network have been proposed to regulate distinct functional domains of AUD development^27–43^. Likewise, several previous studies have focused on the role of Hcrt system activity in regulation of feeding and motivated behaviors, specifically through interactions with mesolimbic dopamine reward pathways^44–48^.

In contrast to interactions with mesolimbic dopamine, the Hcrt system also displays prominent interactions with neuronal stress pathways to drive reinstatement of drug-seeking behavior in rodent models of stress-induced relapse^33,49–51^. These data support a role for the Hcrt system in the withdrawal/negative affective stage of the addiction cycle, consistent with recent human clinical data indicating the potential utility of Hcrt receptor blockers for managing stress reactivity, anxiety, insomnia, and AUD^52–55^. Indeed, Hcrt neurons innervate the extended amygdala^56,57^, including the BNST in particular^58–61^, which forms critical circuits regulating negative reinforcement in AUD through corticotropin-releasing factor (CRF) and other stress neuropeptides^62–65^. The BNST is enriched with steroid hormone receptors, allowing sex-specific modulation of neuronal activity and motivated behaviors^66–68^. For example, one study reported a greater number of CRF neurons in the BNST of female mice, compared to males^69^. Furthermore, female mice display greater excitatory synaptic input onto BNST CRF neurons compared to males, and this sex difference was found to be equalized following alcohol drinking^70^. Likewise, stress-associated activity of the Hcrt system is linked to differential responses in male vs. female rats^71–74^, suggesting that addiction relevant mechanisms of Hcrt/BNST interactions may also differ by sex. Considering that male and female mice display hormone-dependent differences in alcohol consumption and related phenotypes^75–78^, we hypothesized that these effects may be driven in part by sexually divergent mechanisms involving Hcrt signaling effects on BNST physiology.

While actions of the CRF system in the BNST have been heavily implicated in the underlying neurobiology of AUD^10,62–64^, only a limited number of addiction-related studies have examined the role of HcrtR signaling in the BNST. With regard to AUD-relevant behaviors, Ubaldi et al. reported that pharmacological antagonism of HcrtR1 in the BNST blocked neuropeptide S-induced reinstatement of alcohol-seeking^79^; however, Campbell et al. discovered that intra-BNST pharmacological antagonism of HcrtR1 did not alter reinstatement of alcohol-seeking^31^. Both studies were conducted in male rats, which may have limited translation to females given sex differences in alcohol drinking^75^ and the BNST transcriptome^80,81^, including BNST HcrtR1 expression levels^82^. In addition, Hcrt signaling in the BNST is known to physiologically activate BNST neurons and increase anxiety^83^, but it remains unknown which Hcrt receptor subtypes and BNST cell types are involved.

These pharmacological studies highlight the importance of further dissecting how HcrtR signaling differentially regulates alcohol intake not only across brain regions, but also, in a cell type-specific manner within a given brain region. Considering the interactions between Hcrt and stress signaling pathways^84–86^, we hypothesized that selective disruption of HcrtR1 and/or HcrtR2 signaling in CRF neurons may reduce alcohol intake, perhaps through relieving negative affective states. Furthermore, as the BNST is a nexus of neuropeptide communication crucial for negative reinforcement processes in AUD^2,59,87–90^, we hypothesized that the BNST is a key region mediating CRF-specific HcrtR signaling effects on excessive alcohol drinking, withdrawal-heightened anxiety behavior, and neurophysiological excitability changes in response to alcohol.

Here we employed CRF-specific Cre-mediated genetic deletion of each Hcrt receptor alone or in combination^91–93^ and tested alcohol drinking, anxiety, and ex vivo excitability of BNST neurons. We found that CRF-specific HcrtR1 deletion reduced alcohol intake and produced sex-specific effects on BNST excitability and synaptic drive, and that CRF-specific deletion of both HcrtRs had sex-specific effects on reducing alcohol drinking and withdrawal-heightened anxiety, as well as BNST excitability. Overall, our findings support a framework in which HcrtR signaling in CRF neurons interacts with sex differences in BNST physiological excitability to regulate long-term excessive alcohol drinking and the persistence of negative affective behaviors through the time course of withdrawal.

## Methods

### Animals

Adult (8- to 16-week-old) male and female mice on a C57BL/6J (B6) background were housed at constant temperature and humidity, under a regular circadian light-dark cycle (lights-on at 7:00 am, lights-off at 7:00pm). Food and water were available ad libitum except for brief periods during behavioral assays. All experiments were performed in accordance with guidelines described in the US National Institutes of Health Guide for the Care and Use of Laboratory Animals and approved by Stanford University Administrative Panel on Laboratory Animal Care.

*Hcrtr1* floxed and *Hcrtr2* floxed mouse lines were generated by Dr. Anne Vassalli as described^91–93^. To conditionally delete *Hcrtr1* and/or *Hcrtr2*, each floxed line was independently crossed to a CRF-Cre driver line (JAX #012704), generating *Crf+/Cre;Hcrtr1^flox/flox^*, *Crf+/Cre;Hcrtr2^flox/flox^*, and *Crf+/Cre;Hcrtr1^flox/flox^;Hcrtr2^flox/flox^*mutant mice. Cre-negative littermates were used as genetic control groups for each separate single or double KO line. Genotyping for generic Cre and CRF-Cre followed protocols from the Jackson Laboratory. For *Hcrtr1^flox/flox^* and *Hcrtr2^flox/flox^* lines, assays for wild-type, floxed, and recombined alleles were developed based on prior publications^91^, and optimized for automated genotyping (Transnetyx, Inc.).

Mice were bred and weaned at 25 days of age, then same-sex housed 2 to 5 per cage in Innovive cages. At 8 to 16 weeks of age, mice were single housed and transferred to Plexiglass cages in the same room for an additional 7-10 days of acclimation period prior to the initiation of the experiments. During the acclimation period, mice received 24 hours access to two 25-ml glass cylinder bottles with metal sipper tubes (both containing water) on either side of the cage, with food evenly distributed along the cage top.

### Two bottle choice intermittent access alcohol drinking

After the acclimation period and baseline anxiety behavioral assays (described below), individually-housed mice underwent an eight-week two bottle choice alcohol drinking experiment during which they received 24-hour access beginning at 3-4 hours after light onset every Monday, Wednesday, and Friday of the week to two bottles: one containing water, and one containing 10% or 20% v/v EtOH dissolved in water. During the first week, mice had access to 10% EtOH and water; starting from Week 2, mice had access to 20% EtOH and water (Fig. 1A). Mice were weighed every week; fluid levels from each bottle and food weights were recorded on a daily basis to calculate primary variables of interest, as previously described^94,95^.

**Figure 1.**
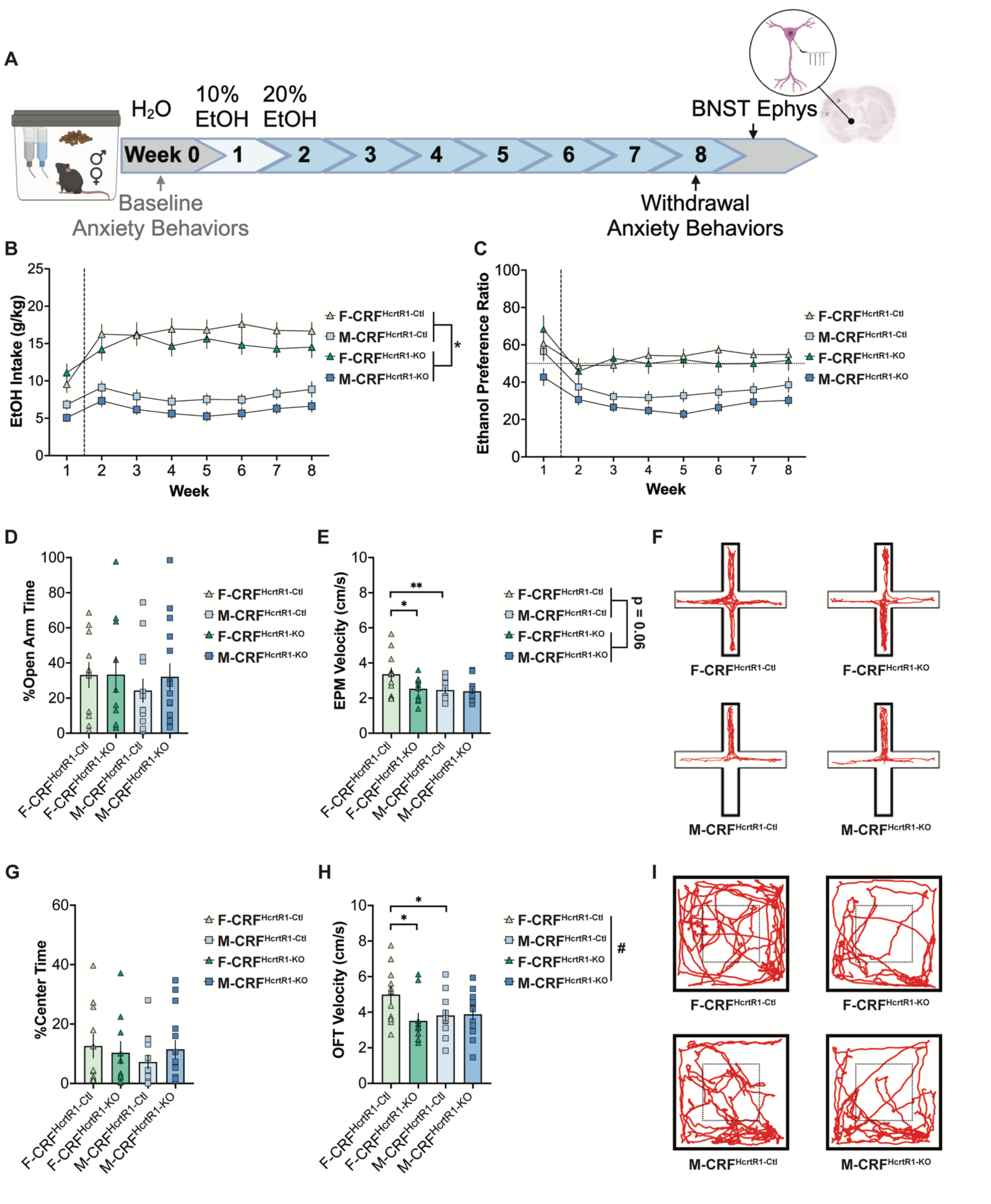
HcrtR1 deletion in CRF neurons reduced alcohol intake. **A,** Timeline for alcohol drinking, behavioral tests, and electrophysiology studies. **B,** Ethanol intake during 8 weeks of two-bottle choice (2BC) intermittent access (IA) from female control (F-CRF^HcrtR1-Ctl^), male control (M-CRF^HcrtR1-Ctl^), female knockout (F-CRF^HcrtR1-KO^), and male knockout (M-CRF^HcrtR1-KO^) mice. Effects of *Genotype p = 0.048, Sex p < 0.0001, Week x Sex p = 0.0010. **C,** Ethanol preference ratio from four groups during 8 weeks of 2BC IA. Effect of Sex p < 0.0001, Week p = 0.0473, Week x Sex p = 0.0003. **D,** Percentage time spent in the open arms in EPM tests during acute abstinence after 8 weeks of 2BC IA. **E,** Velocity during the EPM test across 4 groups. Effect of Sex, p = 0.0289, Genotype p = 0.0555. *F-CRF^HcrtR1-Ctl^ vs F-CRF^HcrtR1-KO^ p = 0.0201, **F-CRF^HcrtR1-Ctl^ vs M-CRF^HcrtR1-Ctl^ p = 0.0080. **F,** Representative motion traces from 4 groups during EPM. **G,** Percentage time spent in the center zone in OFT during acute abstinence after 8 weeks of 2BC IA. **H,** Velocity in the OFT across 4 groups. ^#^Effect of Sex x Genotype, p = 0.0439, Genotype p = 0.0654, *F-CRF^HcrtR1-Ctl^ vs F-CRF^HcrtR1-KO^ p = 0.0119, *F-CRF^HcrtR1-Ctl^ vs M-CRF^HcrtR1-Ctl^ p = 0.0311. **I,** Representative motion traces from 4 groups during OFT. N = 10-15 animals per sex per genotype.

### Anxiety-like behavior testing

Baseline elevated plus maze (EPM) and open field test (OFT) anxiety-like behavioral assays were performed after the acclimation period, prior to alcohol access. Anxiety-like behaviors during acute withdrawal were conducted during the last week of alcohol drinking (Week 8), 5-8 hours after removal of alcohol access. Anxiety-like behaviors during protracted withdrawal were measured by EPM, OFT, and the novelty-suppressed feeding task (NSFT) two weeks after the last access to alcohol. Tests occurred at 8-11 hours after light onset and were separated by at least 24-hour intervals. Mice were video-recorded for 5 min immediately after being placed in the center of the EPM or OFT apparatus. The EPM consisted of a white acrylic apparatus elevated 39 cm off the floor, consisting of two symmetrical open arms and two symmetrical closed arms (each arm 30 cm x 5 cm), with a 5 cm x 5 cm square connecting the four arms in the center. The OFT consisted of a 50 cm x 50 cm white acrylic square chamber, with the center of the arena defined as the inner 25 cm x 25 cm square.

For the NSFT, mice were food deprived for 24 hours prior to the test. A regular chow pellet was placed in the center of the OFT. Mice were video-recorded for 10 min immediately after being placed in the OFT center chamber next to the pellet. Food zone was defined as the inner 12.5 cm x 12.5 cm square. After the 10 min sessions, the same food pellet was weighed immediately and then placed in the cage for an additional 4 hours. Pellets were weighed 2 hours after the behavioral session ended to calculate food intake.

We used eZtrack^96,97^ to quantify EPM, OFT, and NSFT videos for distance traveled and open arm time/center time/food zone time. Distance traveled was converted to mean velocity (cm/s) for data analysis and presentation. For NSFT, latency to investigate food was hand-scored.

### Electrophysiology

Patch-clamp recordings were performed on acute brain slices from both alcohol-naïve and alcohol-experienced mice to assess neuronal excitability and spontaneous synaptic activity. For acute withdrawal groups, slices were collected 3-10 days after last alcohol access; for protracted withdrawal, slices were collected 25-31 days after last alcohol access. 300 μm coronal brain sections through the BNST were collected in ice-cold sucrose-based cutting solution (in mM: 194 sucrose, 20 NaCl, 4.4 KCl, 2 CaCl2, 1 MgCl2, 1.2 NaH2PO4, 26 NaHCO3, and 10 glucose) saturated with carbogen (95% O2/5% CO2). Slices were incubated at 32-34 °C in the aCSF (in mM: 124 NaCl, 4.0 KCl, 1 NaH2PO4, 1.2 MgSO4, 2 CaCl2, 26 NaHCO3, and 10 dextrose, saturated with carbogen) for 30-60 mins. After incubation, the slices were kept at room temperature and were transferred to a recording chamber that was constantly perfused with carbogen-saturated aCSF at a rate of 1-2 ml/min. Recordings were conducted at 28± 2°C. No EtOH was present in the slice bath.

For intrinsic excitability recordings, anterior dorsolateral BNST neurons (including CRF and non-CRF) were visualized with DIC on a Nikon Eclipse FN1 microscope. For synaptic input recordings, only CRF neurons expressing tdTomato in the anterior dorsolateral BNST were visualized with a white LED through a CY3/TRITC filter cube. Whole-cell patch clamp recordings were conducted using 3-5 MΩ micropipettes. Recordings were amplified using an AM Systems amplifier, low-pass filtered at 2 kHz and digitally sampled at 10 kHz using a digitizer (Molecular Devices). Intrinsic excitability experiments used a K-gluconate based internal solution (in mM: 135 K-gluconate, 5 NaCl, 2 MgCl2, 0.6 EGTA, 4 Na2ATP, 0.4 Na2GTP, and 10 HEPES, pH 7.3, 290±5 mOsm). After a gigaseal was formed, neurons were recorded for 1 min to measure on-cell spontaneous firing. After breaking into the whole-cell configuration, neurons were stabilized for 2 mins, then resting membrane potential (RMP) was measured. Neurons with RMP < −40mV were included in the analysis. Ramp current was injected at a rate of 24pA/s to determine action potential threshold. The h-current was measured using a 750ms, −60mV to −100mV voltage step. Evoked spike frequency was measured during 250ms depolarizing current steps ranging from 0-160 pA. Hyperpolarizing current pulses utilized 50 pA steps.

A cesium-based internal solution was used to measure spontaneous synaptic activity (in mM: 135 cesium methanesulfonate, 10 KCl, 1 MgCl2, 0.2 EGTA, 4 MgATP, 0.3 Na2GTP, 20 phosphocreatine, 1mg/ml QX-314, pH 7.3, 290±5 mOsm). Recordings were low-pass filtered at 5 kHz and digitally sampled at 10 kHz. Cells were held at −55mV to assess spontaneous excitatory postsynaptic currents (sEPSCs), and at +10mV to assess spontaneous inhibitory postsynaptic currents (sIPSCs), voltages near the reversal potentials for GABAergic and glutamatergic PSCs, respectively. Input resistance and series resistance were monitored with a 5-mV pulse (10 ms) between epochs, and only recordings that remained stable over the period of data collection were used. Chloride reversal potential for these conditions was calculated as approximately −61.69mV, without correcting for liquid junction potential.

### Fluorescent in situ hybridization (FISH)

Fresh frozen coronal sections containing BNST were processed for RNAscope according to manufacturer’s guidelines. Hybridization was performed using RNAscope Fluorescent Multiplex Kit v1 (Advanced Cell Diagnostics). Transcripts examined were *Hcrtr1* (ACD #466631-C2) and *Crh* (ACD #316091). Slides were coverslipped with mounting medium with DAPI (ab104139) and stored at 4C in the dark until imaging. Images were taken on Zeiss LSM 900 confocal microscope at 20x magnification, processed with ZEN software and FiJi.

### Data Analysis

Statistical tests were performed in GraphPad Prism. Significance threshold was set at p < 0.05. On the figures, significant main effects of single variable are noted by * and ** (p < 0.05, 0.01); significant effects of interaction are noted by # (p < 0.05). Data are expressed as mean + standard error of the mean (SEM). Figures were created with GraphPad Prism, Origin, and BioRender.

#### Ethanol Drinking

EtOH consumption in mL was converted to grams based on density of EtOH concentration and divided by the animal’s body weight to give daily intake scores expressed in grams per kilogram (g/kg). Daily EtOH preference was calculated by dividing EtOH consumption in mL by the total fluid consumption in mL (EtOH consumption + H2O consumption). Data points across three EtOH drinking sessions per week were averaged within each animal to produce a single weekly 24-hour value for EtOH intake (g/kg) and ethanol preference ratio. Each dependent variable was analyzed by a 2×2×7 repeated measures (RM-) ANOVA design with genotype (KO, WT) and sex (male, female) as the between-subjects factors, and week of 20% EtOH intake (weeks 2 - 8 of EtOH intake) as the repeated measure.

#### Anxiety-like Behaviors

Each variable was analyzed by a 2×2 or 2×2×2 RM-ANOVA design with genotype (KO, WT) and sex (male, female) as the between-subjects factors, and withdrawal conditions (acute withdrawal, protracted withdrawal) as the repeated measure. For non-normal data sets, measurements were transformed by natural log or square root and then analyzed by RM-ANOVA. Post-hoc tests were Fisher’s LSD test and Bonferroni correction for multiple comparisons.

#### Electrophysiology

Recordings were analyzed in Clampfit (Molecular Devices). Intrinsic excitability measurements were analyzed by a 2×2 ANOVA design with genotype (KO, WT) and sex (male, female) as the between-subjects factors. For non-normal data sets, measurements were transformed by natural log and then analyzed by RM-ANOVA. Post-hoc tests were Fisher’s LSD test and Bonferroni correction for multiple comparisons. The effects of sex and genotype on spontaneous firing were analyzed via multiple logistic regression.

Spontaneous recordings were filtered at 2kHz with LowPass Elliptic filter and analyzed in the MINI program. Each event was confirmed by the experimenter. At least 150 events were included for each neuron, and the event threshold was set at least 3 times of the noise level.

## Results

### HcrtR1 deletion in CRF neurons reduced alcohol intake

To investigate if disrupting HcrtR signaling in CRF neurons altered alcohol drinking, we created a genetic model by crossing CRF-Cre driver mice with Cre-dependent *Hcrtr1* floxed mice. This selective deletion of HcrtR1 in CRF-expressing neurons produced the *Crf+/Cre;Hcrtr1^flox/flox^* (CRF^HcrtR1-KO^) line. Cre-negative littermates, *Hcrtr1^flox/flox^* (CRF^HcrtR1-Ctl^) served as the control group. Thus, comparing alcohol drinking between CRF^HcrtR1-KO^ and CRF^HcrtR1-Ctl^ mice tested the hypothesis that selective inactivation of HcrtR1 signaling in CRF neurons impacts these behaviors. By performing RNAscope *in situ* hybridization in slices containing dorsal and ventral BNST, we verified that *Hcrtr1* mRNA was selectively lost from CRF neurons but remained present on non-CRF neurons (Fig. S1).

We employed a chronic two-bottle choice (2BC) intermittent access (IA) alcohol drinking paradigm to measure voluntary alcohol consumption over 8 weeks (Fig. 1A). Notably, CRF^HcrtR1-KO^ mice consumed significantly less alcohol than their control counterparts, indicating that targeted disruption of HcrtR1 signaling in CRF neurons reduces alcohol intake (Fig. 1B). As noted in previous studies^75,77,78^, females displayed greater overall alcohol intake and preference compared to males (Fig. 1B-C).

### HcrtR1 deletion in CRF neurons did not significantly impact anxiety-like behavior

Given the roles of CRF and Hcrt signaling in negative reinforcement and hyperkatifeia, we hypothesized that the diminished alcohol consumption in CRF^HcrtR1-KO^ mice could be due to reduced negative affective states during withdrawal. To assess this, we performed the elevated plus maze (EPM) and open field test (OFT) before mice had access to alcohol, and again during acute withdrawal during the eighth week of alcohol drinking (5-8 hrs after cessation of the drinking session). At baseline before alcohol exposure, CRF-specific HcrtR1 deletion did not impact anxiety-like behaviors (Fig. S2). As CRF^HcrtR1-KO^ mice consumed less alcohol, we predicted that CRF-specific HcrtR1 deletion would have an anxiolytic effect, especially during acute withdrawal following week 8 alcohol drinking sessions. However, during acute withdrawal, CRF^HcrtR1-KO^ and control mice also did not significantly differ in anxiety-like behaviors across both tests (Fig. 1D, 1G). Although we did observe trends toward lower overall locomotor activity levels in CRF^HcrtR1-KO^ mice compared to controls (Fig. 1E, 1H), these were unlikely to significantly impair display of anxiety-like behaviors. Together, these findings indicate that CRF-specific HcrtR1 signaling is required for excessive alcohol intake but may not be required for baseline or acute withdrawal-heightened anxiety-like behaviors.

### HcrtR1 deletion in CRF neurons produced sex-specific effects on intrinsic excitability and excitatory drive onto BNST neurons from alcohol-naïve mice

Based on our observation that CRF-specific HcrtR1 deletion reduced alcohol intake, we hypothesized that these effects occurred in part due to altered baseline or alcohol-induced changes in excitability or synaptic drive onto BNST neurons. In dorsolateral BNST neurons from alcohol-naïve mice, we first identified overall sex differences in input resistance (Rm, greater in females) and Ih current (greater in males) (Fig. S3B-C). In addition, we identified a significantly higher actional potential threshold in BNST neurons from male CRF^HcrtR1-KO^ mice compared to male controls (Fig. S3D). These findings suggest that male CRF^HcrtR1-KO^ mice may display decreased alcohol consumption due to altered intrinsic excitability in BNST neurons prior to alcohol drinking. We further tested whether HcrtR1 deletion in CRF neurons could modulate spontaneous excitatory/inhibitory synaptic inputs onto BNST CRF neurons from alcohol-naïve mice. Intriguingly, CRF-specific HcrtR1 deletion increased both the amplitude and frequency of spontaneous excitatory postsynaptic currents (sEPSCs) in females only (Fig. S4A-B). Inhibitory inputs onto BNST CRF neurons were similar across sexes and genotypes (Fig. S4C-D). These findings indicate that, prior to alcohol exposure, CRF-specific HcrtR1 deletion in females can alter the pattern of excitatory drive onto BNST CRF neurons, which may function to limit subsequent alcohol consumption.

### HcrtR1 deletion in CRF neurons produced sex-specific effects on intrinsic excitability in BNST neurons from alcohol-experienced mice

Next, we performed electrophysiological investigations of dorsolateral BNST neurons from CRF^HcrtR1-KO^ and control mice that underwent 8 weeks of alcohol drinking. We found that CRF-specific HcrtR1 deletion blunted the significant sex difference in resting membrane potential (RMP) that was observed among control littermates, with male CRF^HcrtR1-KO^ mice trending toward a lower RMP relative to male controls (Fig. 2A). Notably, we also found that male CRF^HcrtR1-KO^ mice displayed significantly greater current-induced firing rates compared to male controls (Fig. 2F). Overall, these findings indicate that among alcohol-experienced mice, CRF-specific HcrtR1 deletion blunted sex differences in intrinsic excitability and enhanced evoked BNST firing rates specifically in males. This elevated BNST excitability in CRF^HcrtR1-KO^ male mice may be reflective of their reduced alcohol drinking relative controls, which may translate into altered patterns of BNST neuronal activity during protracted withdrawal and under conditions of stress-induced relapse.

**Figure 2.**
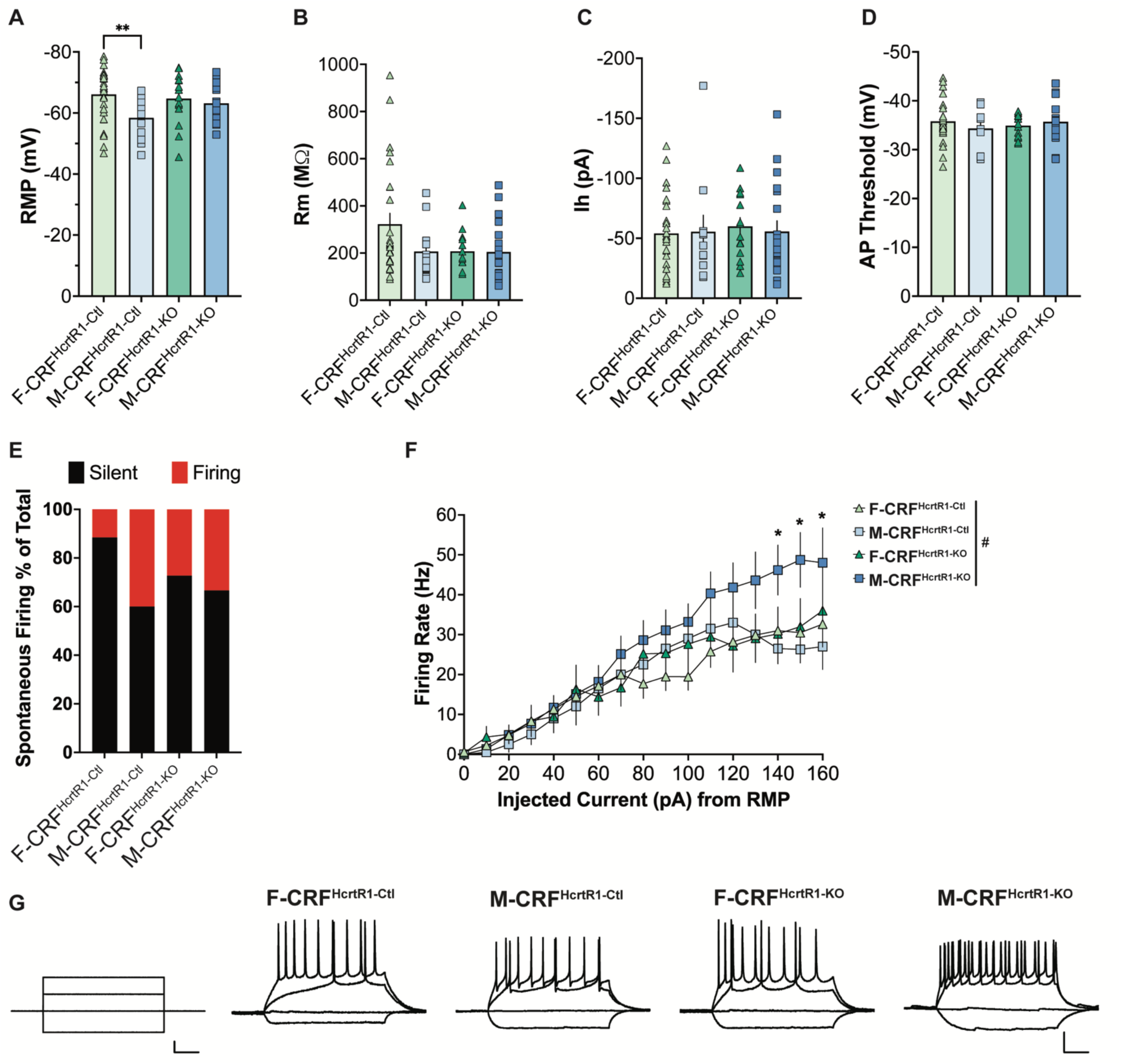
HcrtR1 deletion in CRF neurons increased BNST intrinsic excitability after chronic alcohol drinking in a sex-specific manner. **A - D,** Resting membrane potential (RMP), input resistance (Rm), h current (Ih), and action potential (AP) threshold of BNST neurons in four groups of mice after 8 weeks 2BC IA alcohol drinking. **A,** RMP Effect of Sex p = 0.0171, **F-CRF^HcrtR1-Ctl^ vs M-CRF^HcrtR1-Ctl^ p = 0.0047, M-CRF^HcrtR1-Ctl^ vs M-CRF^HcrtR1-KO^ p = 0.0934. **E,** Percentage of spontaneously firing BNST neurons was similar across groups. **F,** CRF-specific deletion of HcrtR1 in males led to higher evoked firing rate of BNST neurons after chronic alcohol drinking. Effects of ^#^Sex x Genotype p = 0.0086, Sex p = 0.0046, Genotype p = 0.0001, *M-CRF^HcrtR1-Ctl^ vs M-CRF^HcrtR1-KO^ p = 0.0392, 0.0188, 0.0192. **G,** Current steps (left) and representative evoked firing recordings for each sex/genotype. Scale bars, 50ms, 50pA, 20mV. N = 11-28 cells/2-4 animals per sex per genotype.

### HcrtR2 deletion in CRF neurons produced sex-specific effects on anxiety behavior

We next explored whether CRF-specific HcrtR2 deletion could reduce alcohol drinking similarly to CRF-specific HcrtR1 deletion. We utilized the same strategy to establish the CRF^HcrtR2-KO^ and CRF^HcrtR2-Ctl^ groups, which underwent 8 weeks of 2BC IA drinking and anxiety-like behavior tests. Unlike HcrtR1 deletion, CRF-specific HcrtR2 deletion did not significantly impact alcohol intake or preference (Fig. 3A-B). In addition, CRF^HcrtR2-KO^ and CRF^HcrtR2-Ctl^ mice performed similarly in the EPM (Fig. 3C). However, in the OFT, male CRF^HcrtR2-KO^ mice displayed decreased anxiety-like behavior compared to female CRF^HcrtR2-KO^ mice, a sex difference that was not present among controls (Fig. 3F). This male-biased anxiolytic effect of CRF-specific HcrtR2 deletion was also observed during baseline anxiety tests, in which male CRF^HcrtR2-KO^ mice spent more time in the center of the OFT compared to male controls (Fig. S5C). Taken together, these results indicate that conditional deletion of HcrtR2 in CRF neurons did not significantly change alcohol drinking but did decrease anxiety-like behaviors among males.

**Figure 3.**
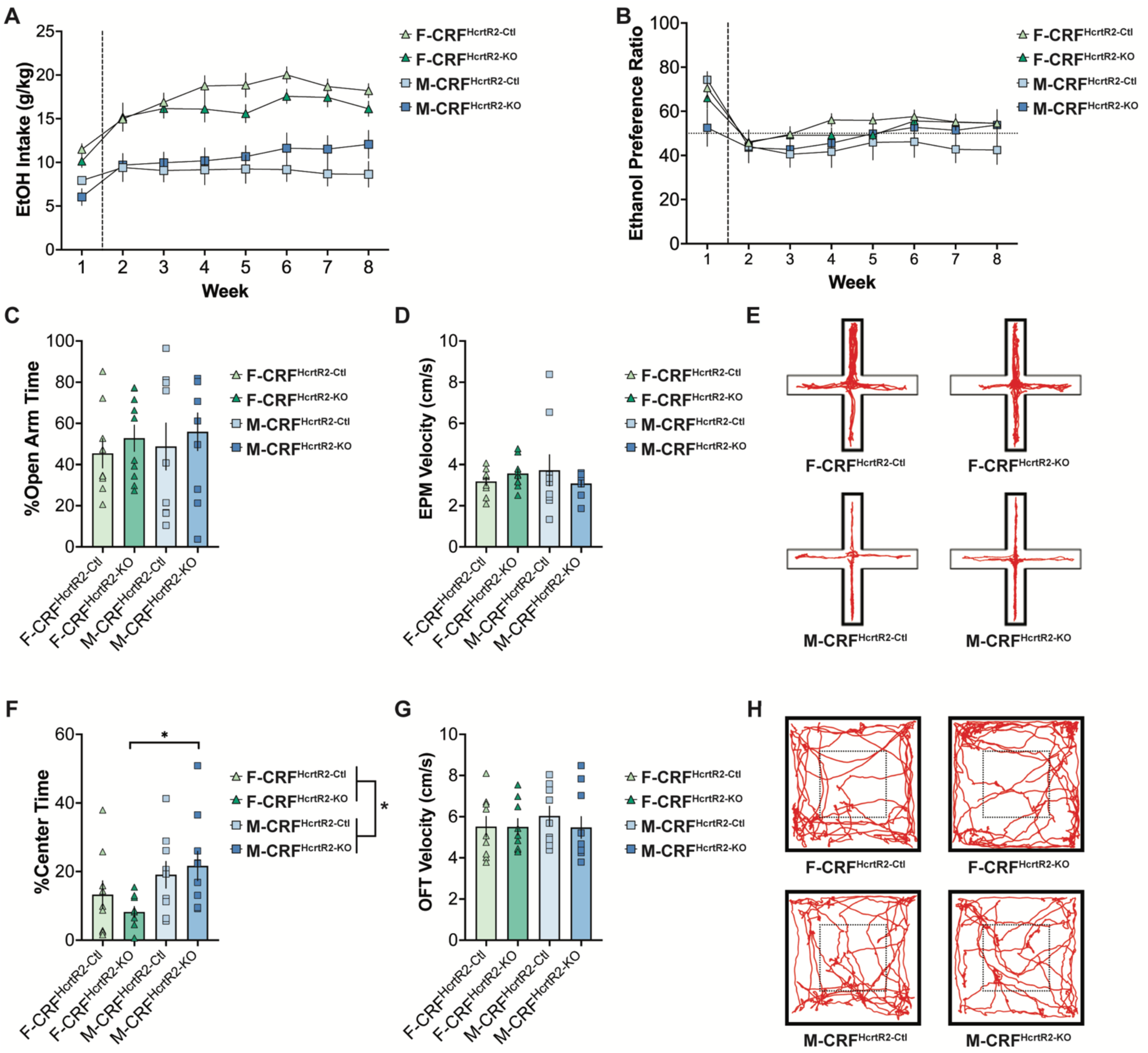
HcrtR2 deletion in CRF neurons produced sex-specific effects on anxiety behavior. **A,** Ethanol intake during 8 weeks of 2BC IA paradigm from female control (F-CRF^HcrtR2-Ctl^), male control (M-CRF^HcrtR2-Ctl^), female knockout (F-CRF^HcrtR2-KO^), and male knockout (M-CRF^HcrtR2-KO^) mice. Effect of Sex p < 0.0001, Week p = 0.0003. **B,** Ethanol preference ratio from four groups during 8 weeks of 2BC IA. Effect of Week p < 0.0001. **C,** Open arm time in EPM tests during acute withdrawal after 8 weeks of 2BC IA. **D,** Velocity during the EPM test across 4 groups. **E,** Representative motion traces from 4 groups during EPM. **F,** Center zone time in OFT during acute withdrawal after 8 weeks of 2BC IA. *Effect of Sex p = 0.0130, *F-CRF^HcrtR2-KO^ vs M-CRF^HcrtR2-KO^ p = 0.0130. **G,** Velocity in the OFT across 4 groups. **H,** Representative motion traces from 4 groups during OFT. N = 9-10 animals per sex per genotype.

### HcrtR2 deletion in CRF neurons produced sex-specific and alcohol experience-dependent effects on BNST excitability

We also characterized the effects of CRF-specific HcrtR2 deletion on BNST excitability. In alcohol-naïve experiments, CRF^HcrtR2-KO^ male mice displayed significantly enhanced current-evoked BNST firing rates (Fig. S6F), reminiscent of HcrtR1 deletion effects on male-specific enhanced firing rates. Otherwise, BNST neurons from female CRF^HcrtR2-KO^ mice exhibited significantly greater input resistance relative to control females (Fig. S6B), and we observed an overall sex difference in AP threshold (Fig. S6D). In BNST neurons from alcohol-experienced mice, female CRF^HcrtR2-KO^ mice exhibited significantly greater current-evoked firing rates compared to female controls (Fig. S6L), despite significantly lower likelihood of spontaneous firing in BNST neurons from CRF^HcrtR2-KO^ mice overall (Fig. S6K). Similar to effects in CRF^HcrtR1-KO^ mice, these data indicate that CRF-specific HcrtR2 deletion enhances current-evoked BNST firing in a pattern that differs between sexes and depends on prior alcohol drinking experience.

### Double deletion of both HcrtRs in CRF neurons produced sex-specific changes in alcohol drinking

Differential effects of CRF-specific HcrtR1 or HcrtR2 deletion on alcohol drinking indicated that these strategies may target separate mechanisms of the Hcrt system’s contributions to AUD processes. Thus, to thoroughly understand the effects of disrupting Hcrt signaling in CRF neurons, we established a double HcrtR KO line (*Crf+/Cre;Hcrtr1^flox/flox^;Hcrtr2^flox/flox^*, CRF^HcrtR1&R2-KO^) and compared them to littermate controls (CRF^HcrtR1&R2-Ctl^) using the 2BC IA protocols. We hypothesized that CRF^HcrtR1&R2-KO^ mice would display reduced alcohol consumption similar to CRF-specific HcrtR1 deletion. Intriguingly, we observed that male CRF^HcrtR1&R2-KO^ mice did indeed display slightly lower levels of alcohol intake and preference, although female CRF^HcrtR1&R2-KO^ mice showed slightly higher levels of alcohol intake and preference (Fig. 4B-C; genotype x sex p = 0.011, p = 0.045).

**Figure 4.**
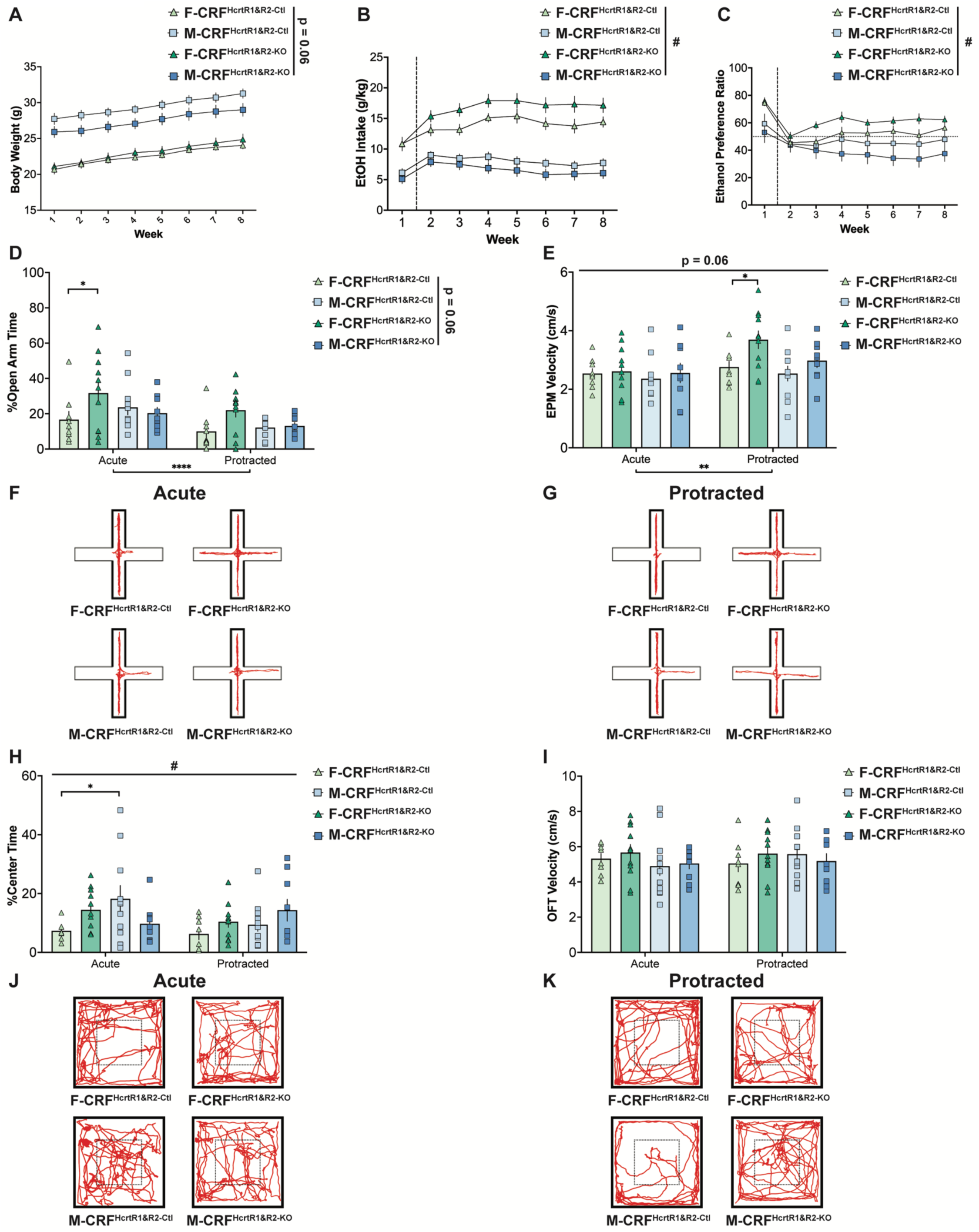
Double deletion of both HcrtRs in CRF neurons had sex-specific impact on alcohol intake and anxiolytic behavioral effects. **A,** Body weight during 8 weeks of 2BC IA paradigm from female control (F-CRF^HcrtR1&R2-Ctl^), male control (M-CRF^HcrtR1&R2-Ctl^), female knockout (F-CRF^HcrtR1&R2-KO^), and male knockout (M-CRF^HcrtR1&R2-KO^) mice. Effect of Sex x Genotype, p = 0.0595, Sex p < 0.0001, Week p < 0.0001. **B,** Ethanol intake from four groups during 8 weeks of 2BC IA. Effects of ^#^Sex x Genotype p = 0.0108, Week x Sex p < 0.0001, Sex p < 0.0001. **C,** Ethanol preference ratio from four groups. Effects of ^#^Sex x Genotype p = 0.0448, Week x Sex p < 0.0001, Sex p = 0.0012, Week p = 0.0352. **D,** Open arm time in EPM tests during acute abstinence (Acute) and after 2 weeks of protracted withdrawal (Protracted) after 2BC IA. Effects of ****Withdrawal Time p < 0.0001, Sex x Genotype p = 0.0581, *Acute F-CRF^HcrtR1&R2-Ctl^ vs F-CRF^HcrtR1&R2-KO^ p = 0.0488. **E,** Velocity during the EPM test across 4 groups. Effects of **Withdrawal Time p = 0.0018, Withdrawal x Genotype p = 0.0582, Genotype p = 0.0617. Protracted *F-CRF^HcrtR1&R2-Ctl^ vs F-CRF^HcrtR1&R2-KO^ p = 0.0476. **F-G,** Representative motion traces from 4 groups in EPM during acute (F) and protracted (G) withdrawal. **H,** Center zone time in OFT during acute abstinence and protracted withdrawal. ^#^Withdrawal Time x Sex x Genotype p = 0.0288, Sex x Genotype p = 0.0945. *Acute F-CRF^HcrtR1&R2-Ctl^ vs M-CRF^HcrtR1&R2-Ctl^ p = 0.0352 (after Bonferroni correction) **I,** Velocity in the OFT across 4 groups. **J-K,** Representative motion traces from 4 groups in OFT during acute (J) and protracted (K) withdrawal. N = 9-11 animals per sex per genotype.

### Double deletion of both HcrtRs in CRF neurons generated anxiolytic behavioral effects

The altered patterns of alcohol drinking in CRF^HcrtR1&R2-KO^ mice hinted at additional neurobehavioral effects as a result of double HcrtR KO. To examine affective behavior at protracted time points following alcohol drinking, we included additional anxiety-like behavioral tests after two weeks of protracted withdrawal following the 8-week 2BC IA paradigm. In baseline behavioral tests, double HcrtR KO did not significantly impact anxiety-like behavior or locomotor activity (Fig. S7). Following alcohol drinking, however, we identified a strong trend toward reduced anxiety in female double KO mice in the EPM, observed across multiple withdrawal timepoints (Fig. 4D, genotype x sex, p = 0.06). This sexually dimorphic anxiolytic effect was more pronounced in the OFT (Fig. 4H, genotype x sex x withdrawal, p = 0.029). We further examined this anxiolytic effect using the novelty suppressed feeding test (NSFT) during protracted withdrawal and confirmed an anxiolytic phenotype, as double KO mice spent significantly more time in the food zone than controls (Fig. S8C, genotype effect p = 0.01). Taken together, these results support our hypothesis that deletion of HcrtRs from CRF neurons via double KO reduced anxiety-like behaviors during alcohol withdrawal.

### Double deletion of both HcrtRs in CRF neurons decreased BNST excitability in protracted withdrawal

Finally, as double HcrtR KO produced anxiolytic effects particularly during protracted withdrawal (Fig. 4H, S8C), we predicted that BNST excitability may be decreased in double HcrtR KO mice after protracted withdrawal. Indeed, we observed significantly reduced evoked firing rate in double HcrtR KO mice compared to controls (Fig. 5F, p = 0.004). We also identified overall sex differences in both spontaneous and evoked firing rates (p = 0.008, p = 0.054). These results indicate that, in contrast to sex-dependent effects of CRF-specific deletion of individual HcrtRs during acute withdrawal, double HcrtR KO during protracted withdrawal led to reduced (rather than enhanced) excitability in BNST neurons from alcohol-experienced mice of both sexes.

**Figure 5.**
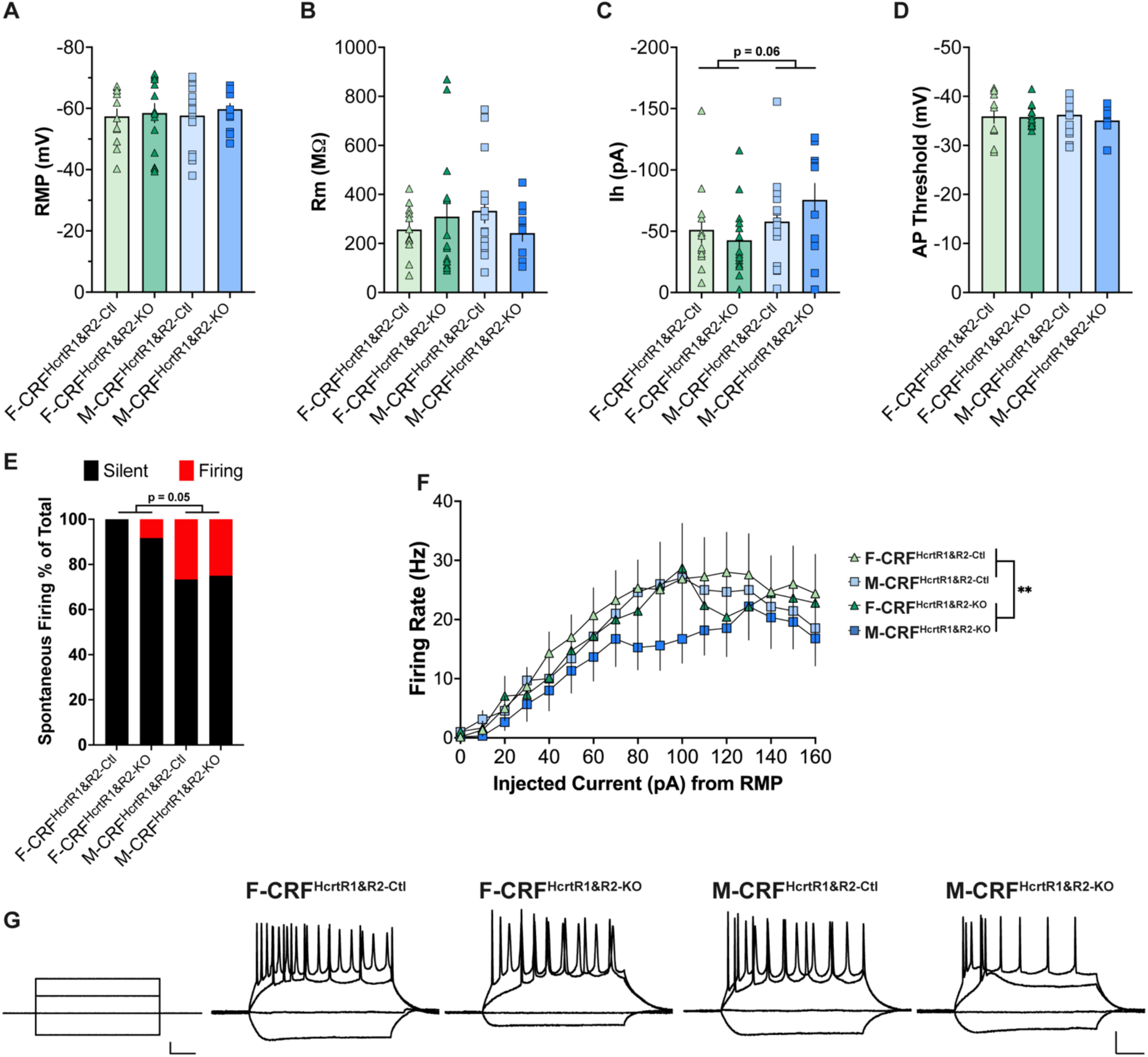
Double deletion of both HcrtRs in CRF neurons modulated BNST excitability during protracted withdrawal. **A-D,** RMP, Rm, Ih, AP threshold were similar across female control (F-CRF^HcrtR1&R2-Ctl^), male control (M-CRF^HcrtR1&R2-Ctl^), female knockout (F-CRF^HcrtR1&R2-KO^), and male knockout mice (M-CRF^HcrtR1&R2-KO^) during protracted withdrawal. **C,** Ih current Effect of Sex p = 0.0565. **E,** Percentage of spontaneously firing in the BNST trended toward higher in males after protracted withdrawal. Effect of Sex p = 0.0543. **F,** CRF-specific deletion of HcrtR1 and HcrtR2 led to lower evoked firing rate of BNST neurons during protracted withdrawal. Effects of **Genotype p = 0.0044, Sex p = 0.0077. **G,** Current steps (left) and representative evoked firing recordings for each sex/genotype. Scale bars, 50ms, 50pA, 20mV. N = 12-16 cells/2 animals per sex per genotype.

## Discussion

Here we successfully identified differential roles for HcrtR1 and HcrtR2 signaling that link their contributions to excessive alcohol drinking, negative affective behaviors, and BNST neuronal excitability. These findings demonstrate importance by bridging two classical stress-arousal systems (Hcrt and CRF) that had not previously been shown to directly interact to regulate addiction-like behaviors in a sex-specific manner. Furthermore, our data are important for revealing that these systems-wide interactions are also associated with sex-specific neurophysiological changes within the BNST of the extended amygdala. Specifically, we have found that CRF-specific HcrtR1 deletion reduced alcohol intake and produced sex-specific enhancements in BNST excitability and synaptic drive (Figs. 1, 2, S4). We went on to show that CRF-specific HcrtR2 deletion produced a male-specific decrease in baseline anxiety-like behavior and enhanced BNST excitability (Fig. 3, S5, S6). Finally, we found that CRF-specific deletion of both HcrtRs reduced alcohol drinking in males as well as dampened anxiety behaviors and BNST excitability in both sexes (Figs. 4, 5, S8).

Recent studies from the alcohol field investigated the role of the Hcrt arousal system in AUD-relevant behaviors, historically focusing on pharmacological approaches^28–38,40–42,79,98–100^. In tandem, contemporary theories of AUD have highlighted the roles of withdrawal and negative affect in perpetuating the addiction cycle^1–3,5,6^, emphasizing the relevance of the extended amygdala and CRF stress neuropeptide system^8,10–13,62,63^. Given prior reports supporting the idea that Hcrt signaling may alter CRF neuronal activity and CRF release^7,47,84–86^, we hypothesized that Hcrt signaling in CRF neurons contributes to the link between substance use and the stress response in a receptor subtype-specific manner. In addition, based on known sex differences in BNST control of motivated behavior^66–69,80,81,101^, sex differences in Hcrt-mediated stress responses^71–74^, and evidence that BNST CRF neurons regulate short-term binge alcohol drinking and anxiety^63,64,70^, we further hypothesized that HcrtR signaling in CRF neurons may differentially modulate behavior and physiological between females and males.

Our studies set out to build on previous finding in several notables ways: 1) to determine whether HcrtR signaling specifically in CRF neurons is required for excessive alcohol drinking and withdrawal-heightened anxiety, 2) to differentiate the roles of HcrtR1 vs. HcrtR2 signaling in these effects, 3) to identify differences in these mechanisms between females and males, and 4) to assess whether such impacts are associated with changes in neuronal excitability specifically within the BNST.

Early anatomical studies of the Hcrt system identified axonal projections innervating the BNST^23^, consistent with documented expression of both *Hcrtr1* and *Hcrtr2* mRNAs in this region^25,26,102^. Later reports suggested the functionality of this Hcrt→BNST pathway, given that Hcrt application to BNST neurons in slice produced physiological spiking activity, and Hcrt microinjections to the BNST increased anxiety behavior and decreased social interaction in mice^83^. Additional findings indicated that these interactions were likely reciprocal. For example, BNST inputs to the Hcrt-LH field were cFos-activated in proportion to the degree of cocaine-induced conditioned place preference^103^. Furthermore, modified rabies tracing, electrophysiological connectivity, and optogenetic behavioral experiments demonstrated that CRF-BNST neurons directly synapse onto Hcrt neurons and drive avoidance behavior via outputs to the LH^104^. Consistent with this framework indicating that reciprocal Hcrt/BNST interactions likely drive maladaptive behaviors underlying negative affect during withdrawal in addiction^10,59,105,106^, blockade of HcrtR1 in the BNST attenuated reinstatement of alcohol-seeking in male rats^79^. Despite these encouraging lines of evidence, open questions remained, including which BNST cell types were involved in Hcrt’s effects on the BNST, whether these effects differed between males and females, and whether these effects were related to long-term excessive 2BC alcohol drinking, withdrawal-associated anxiety, or neurophysiological changes within the BNST.

In our primary set of findings, CRF-specific HcrtR1 deletion significantly reduced alcohol intake in both sexes (Fig. 1B). Given this reduction in alcohol drinking, we predicted that this could be related to anxiety behavior in two ways: 1) decreased anxiety behavior at baseline could protect against the motivation to consume alcohol, and/or 2) decreased alcohol consumption could lead to decreased anxiety during acute withdrawal. Surprisingly, in CRF^HcrtR1-KO^ mice, our data did not support these predictions, as genotypes did not show significant anxiety behavior differences in either the EPM or OFT (Fig. 1D, 1G). It is possible that the effects of CRF-specific HcrtR1 deletion on withdrawal-enhanced anxiety predominate at protracted rather than acute time points, and thus would lead to reduced anxiety only during protracted withdrawal (multiple weeks following drinking). Alternatively, both HcrtR1 and HcrtR2 signaling in CRF neurons may promote anxiety-like behaviors in distinct ways. Consistent with this hypothesis, CRF^HcrtR2-KO^ male mice displayed evidence of decreased anxiety at baseline (Fig. S5A,C). Furthermore, while male CRF^HcrtR1&R2-KO^ mice exhibited reduced alcohol intake and preference, their anxiolytic behavioral effects were more prominent during withdrawal rather than at baseline (Fig. 4D,H, S7A,C, S8C). Overall, our double KO approach generated evidence demonstrating that coordinated signaling of both HcrtR subtypes within CRF neurons is likely required for high levels of alcohol drinking, as well as for withdrawal-heightened anxiety behaviors.

For the electrophysiological assessments of intrinsic excitability in alcohol-naïve vs. alcohol-experienced mice across HcrtR KO lines, our datasets reflect the general population of anterior dorsolateral BNST neurons including non-CRF expressing neurons (as we had limited ability to accurately visualize GFP-tagged CRF-HcrtR KO neurons). Deletion of HcrtR1 from CRF neurons notably enhanced current-induced firing rates in alcohol-experienced males (Fig. 2F). The simplest explanation for this finding is that CRF-specific loss of HcrtR1 in males increases BNST excitability, which results in decreased alcohol consumption. However, given that HcrtR1 signaling is primarily Gq/Gs-coupled, and that greater BNST excitability is generally theorized to be associated with enhanced rather than reduced alcohol drinking, other possibilities remain. For example, our observation of enhanced BNST excitability may be an effect (rather than a cause) of lower overall levels of alcohol drinking in male CRF^HcrtR1-KO^ mice, as prior alcohol drinking may have more efficiently suppressed current-induced firing in BNST neurons from high-drinking male control mice. Alternatively, loss of stimulatory HcrtR1 signaling from CRF cells may have had overall disinhibitory effects on spiking activity in recorded cells, given that the BNST includes subpopulations of local inhibitory interneurons (including some CRF-positive BNST neurons). Finally, it remains possible that compensatory upregulation of HcrtR2 may have contributed to enhanced spiking rates in BNST neurons from male CRF^HcrtR1-KO^ mice. Similarly, our finding that deletion of HcrtR2 from CRF neurons enhanced current-induced spiking in BNST neurons from female mice (Fig. S6L) indicates that compensatory upregulation of HcrtR1 may be driving this analogous effect. Consistent with these hypotheses, our observation of significantly reduced current-induced firing in BNST neurons from CRF^HcrtR1&R2-KO^ mice (Fig. 5F) supports our overall conclusion that HcrtR1 and HcrtR2 each have contributions to BNST excitability that may compensate in each other’s absence. Intriguingly, a previous investigation focused on dopamine neurons also found behavioral and physiological effects of double HcrtR deletion that were somewhat opposed to single deletion of HcrtR2 alone^91^.

In our experiments that characterized effects of CRF^HcrtR1^ deletion on spontaneous excitatory/inhibitory drive, we crossed CRF^HcrtR1-KO^ mice with Cre-dependent tomato reporter mice to guarantee high-accuracy targeting of CRF-positive BNST neurons for synaptic assessments. We found that deletion of HcrtR1 from CRF neurons increased excitatory drive onto BNST CRF neurons, specifically in alcohol-naïve females (Fig. S4A-B). This indicates that under normal physiological conditions, HcrtR1 signaling may constrain excitatory transmission onto BNST CRF neurons. However, this possibility may be unlikely because Gq/Gs-coupled HcrtR1 signaling would be more likely to accentuate, rather than constrain, excitatory inputs. Alternatively, loss of HcrtR1 signaling in non-BNST CRF neurons (for example, in central amygdala, paraventricular thalamus, or nucleus accumbens) may lead to enhanced spontaneous excitatory drive onto BNST CRF neurons through an indirect chain of events. Overall, these datasets demonstrate that CRF-specific deletion of HcrtR1 signaling similarly increased excitation in BNST neurons from both male and female mice. Therefore, this manipulation can increase BNST excitability through divergent sex-specific mechanisms that may vary depending on cell type and prior alcohol drinking history.

In summary, our studies confirmed that HcrtR1 signaling is critical for modulating alcohol intake, and that both HcrtR1 and HcrtR2 contribute to anxiety-like behaviors in a sex-dependent and withdrawal-dependent manner. The differential and synergistic contribution of both HcrtRs, which is in part mediated by BNST circuit excitability, could be explained by their distinct expression patterns across sexes and brain regions. For example, males show higher HcrtR1 expression than females in the BNST^82^. Furthermore, HcrtR1 is more enriched in the locus coeruleus, while HcrtR2 is more abundant in the arcuate nucleus, lateral amygdala, lateral hypothalamus, medial septum, cortex, preoptic area, and tuberomammillary nuclei of the hypothalamus^107,108^. Our studies warrant further sex-specific and circuit-specific investigation of HcrtR signaling at different stages of AUD and indicate that therapeutic approaches targeting the Hcrt system may have beneficial effects through modulation of activity in the BNST and CRF neurons that contribute to negative affective responses during withdrawal.

## Acknowledgments

We thank Valentina Martinez Damonte, Chelsie L. Brewer, Brittany J. Bush, Joseph R. Knoedler, Malia A. Belnap, Leigh C. Walker, Luis de Lecea, and all members of Giardino and Kauer laboratories for their contributions. Funding from NIH/NIAAA K99/R00 AA025677, Whitehall Foundation, Brain Research Foundation (W.J.G.) and NIH/NIDA R01 DA011289 (J.A.K.).

