## Supplemental Figures 1-8 and Legends for "Sex-Specific Roles of Hypocretin Receptor Signaling in CRF Neurons on Alcohol Drinking, Anxiety, and BNST Neuronal Excitability"

### Supplementary Figure 1

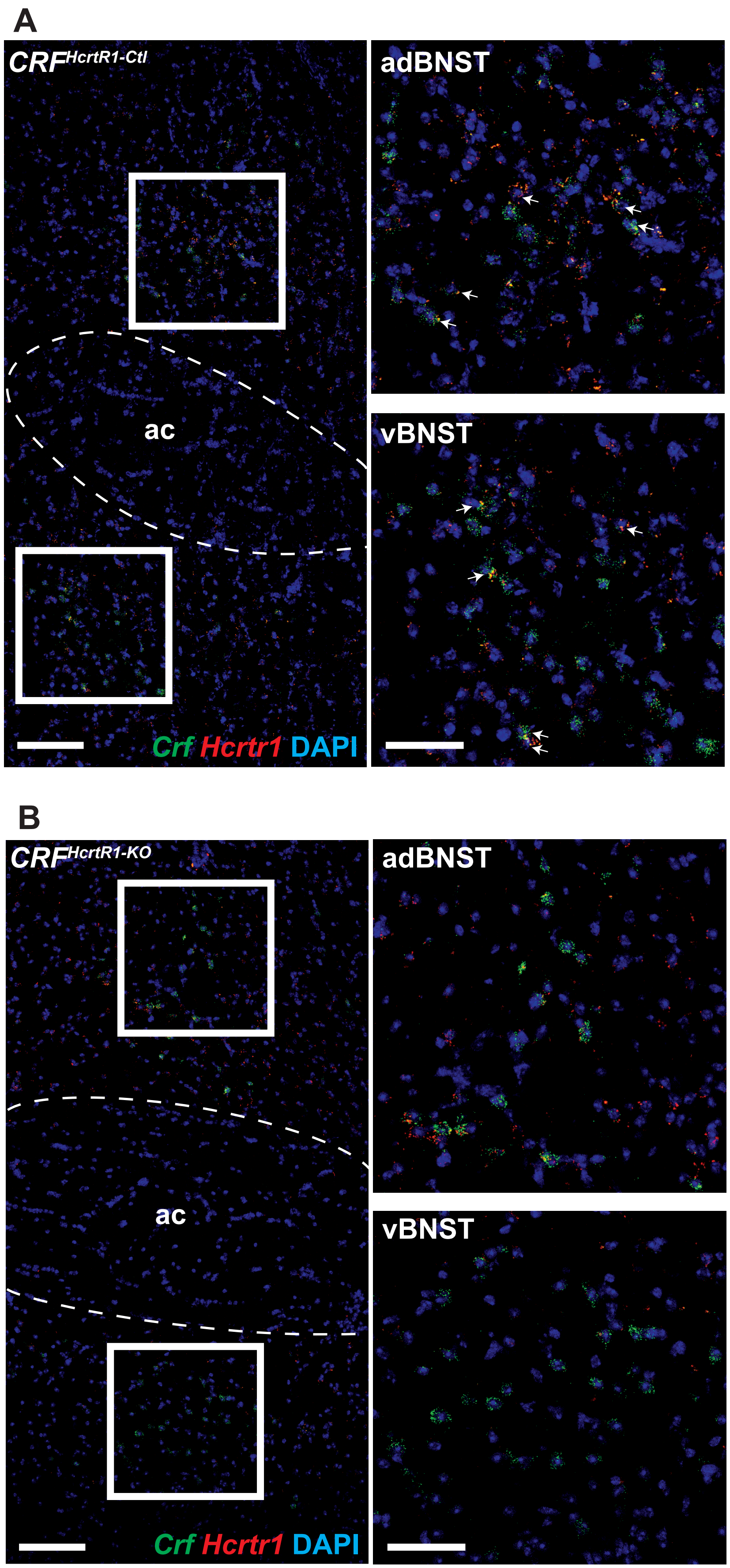

**Figure S1. Fluorescent *in situ* hybridization confirmed selective deletion of *Hcrtr1* expression in CRF neurons.**

**A,** Representative images of RNAscope in the BNST of a control ($CRF^{HcrtR1\text{-}Ctl}$) animal using the *Crf/Crh* probe (green), *Hcrtr1* probe (red), and DAPI (blue). White arrows, cells co-expressed *Crh* and *Hcrtr1*. **B,** Representative images in the BNST of a conditional KO animal ($CRF^{HcrtR1\text{-}KO}$) tested with the same probes but exhibited an absence of co-labelled neurons. Scale bars, 100um (left panels), 50um (right panels).

### Supplementary Figure 2

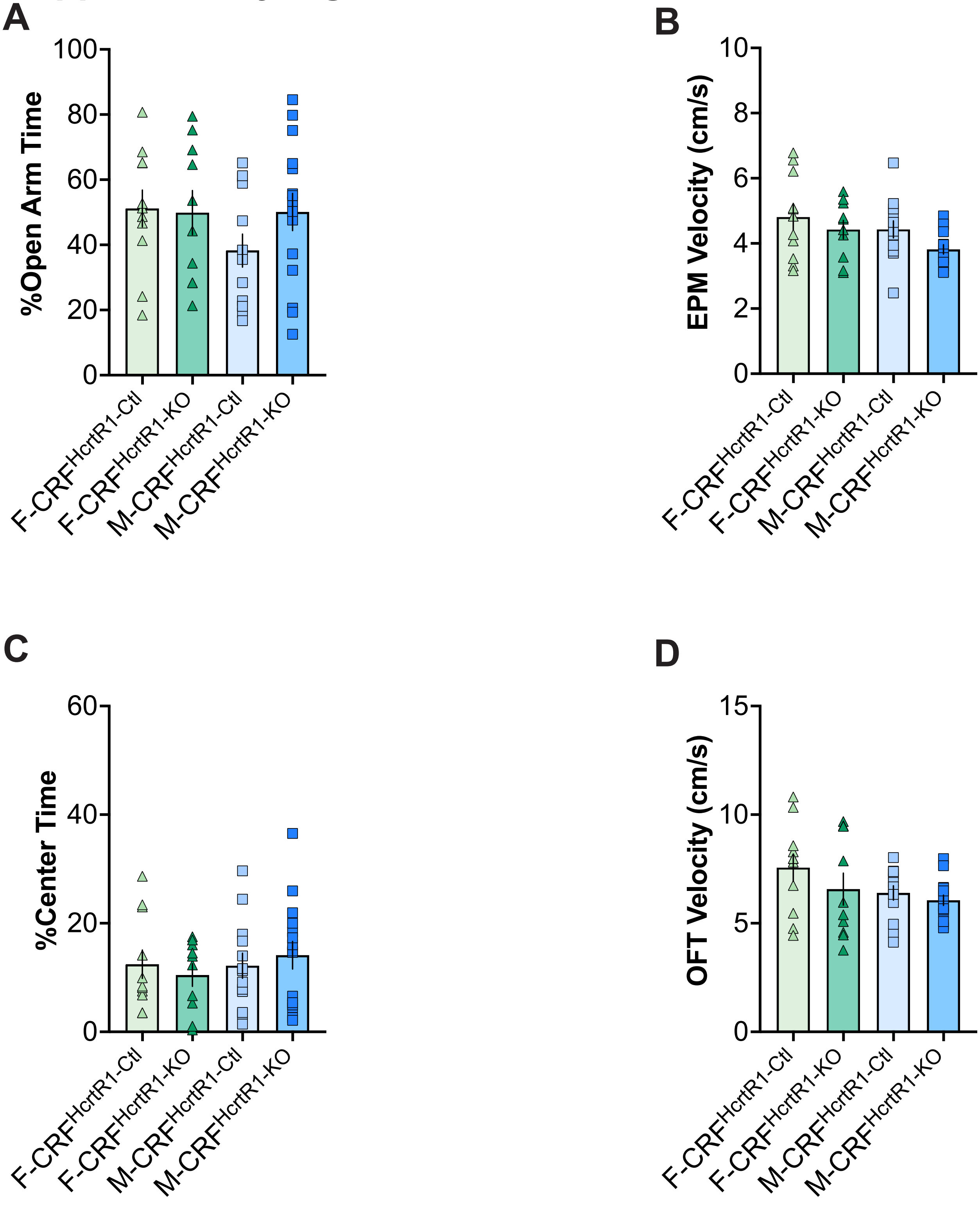

**Figure S2. HcrtR1 deletion in CRF neurons did not significantly impact baseline anxiety-like behavior.**

**A,** Percentage time spent in the open arms in EPM tests from alcohol-naive female control (F-CRF$^{HcrtR1\text{-}Ctl}$), male control (M-CRF$^{HcrtR1\text{-}Ctl}$), female knockout (F-CRF$^{HcrtR1\text{-}KO}$), and male knockout (M-CRF$^{HcrtR1\text{-}KO}$) mice. **B,** Velocity during the EPM test across 4 groups. **C,** Percentage time spent in the center zone in OFT from four groups of alcohol-naive mice. **D,** Velocity in the OFT across 4 groups. N = 10-15 animals per sex per genotype.

### Supplementary Figure 3

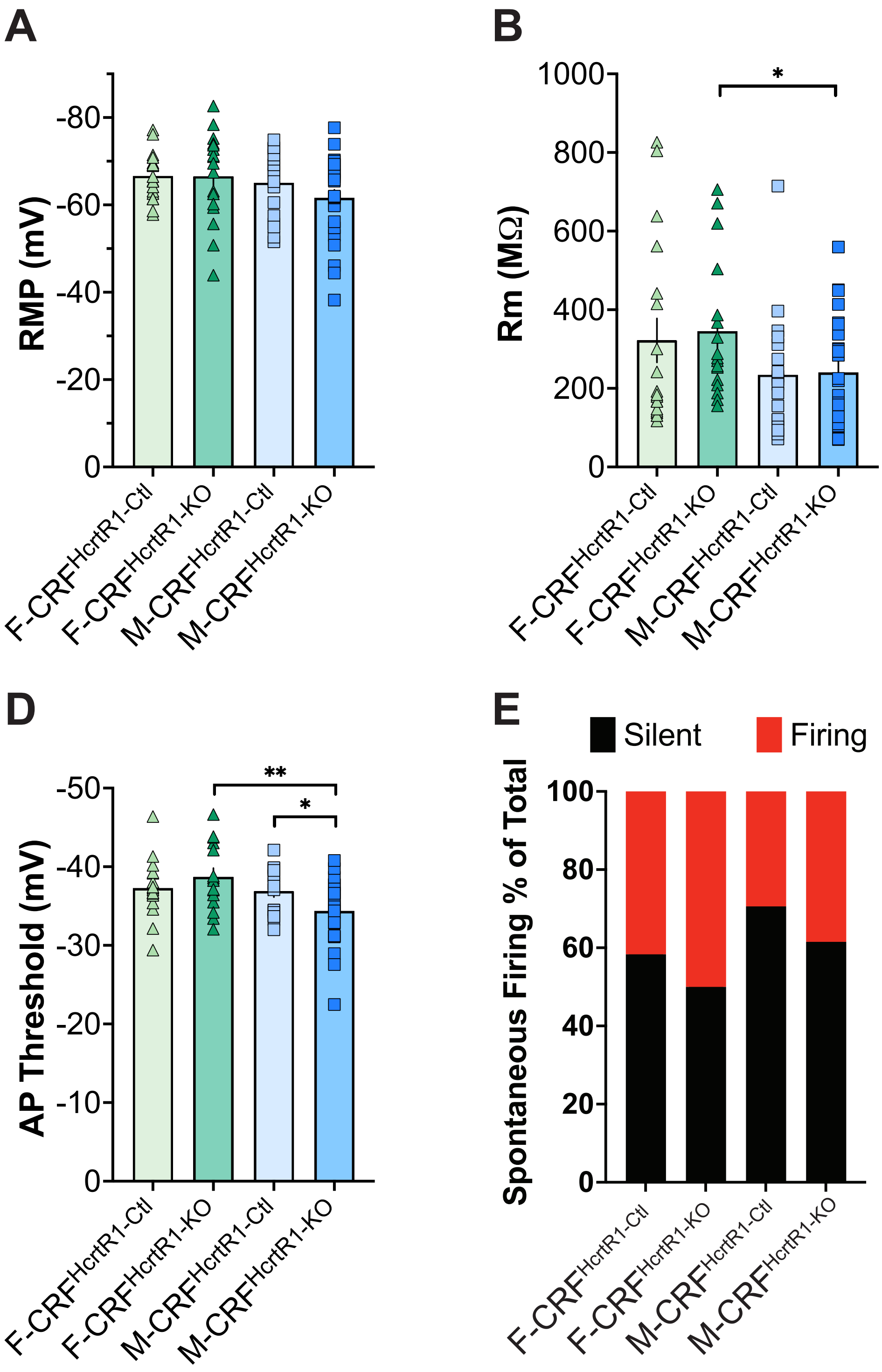

**Figure S3. Alcohol-naïve mice display sex differences in BNST intrinsic excitability.**

**A,** RMP of BNST neurons in alcohol naive female control (F-$CRF^{HcrtR1\text{-}Ctl}$), male control (M-$CRF^{HcrtR1\text{-}Ctl}$), female knockout (F-$CRF^{HcrtR1\text{-}KO}$), and male knockout (M-$CRF^{HcrtR1\text{-}KO}$) mice. **B,** Rm of BNST neurons differed between males and females ($p = 0.0127$). *F-$CRF^{HcrtR1\text{-}KO}$ vs. M-$CRF^{HcrtR1\text{-}KO}$ $p = 0.0255$. **C,** Ih also exhibited a sex difference ($p = 0.0106$). *F-$CRF^{HcrtR1\text{-}KO}$ vs. M-$CRF^{HcrtR1\text{-}KO}$ $p = 0.0186$. **D,** HcrtR1 deletion affected AP threshold of BNST neurons in a sex-dependent manner. Effect of Sex x Genotype, $p = 0.0350$, Sex $p = 0.0113$, **F-$CRF^{HcrtR1\text{-}KO}$ vs M-$CRF^{HcrtR1\text{-}KO}$ $p = 0.0010$, *M-$CRF^{HcrtR1\text{-}Ctl}$ vs M-$CRF^{HcrtR1\text{-}KO}$ $p = 0.0433$. **E,** Percentage of spontaneously firing BNST neurons were similar across groups. N = 15-26 cells/3 animals per sex per genotype.

### Supplementary Figure 4

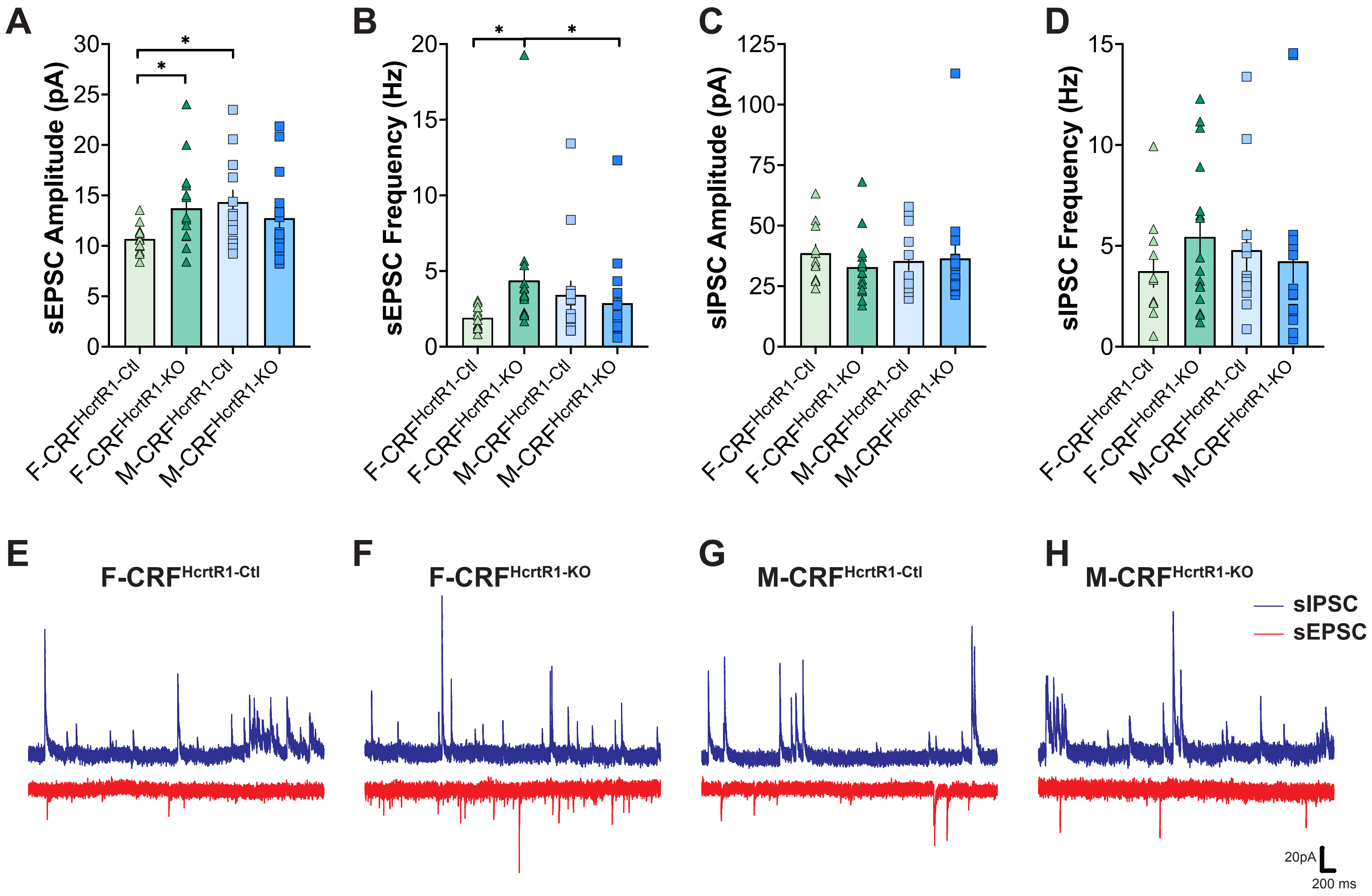

**Figure S4. HcrtR1 deletion in CRF neurons in alcohol-naïve mice selectively increased excitatory synaptic inputs onto BNST CRF neurons in females.**

**A,** Amplitude of spontaneous excitatory postsynaptic currents (sEPSC). Effect of Sex x Genotype p = 0.0262, *F-CRF$^{HcrtR1\text{-}Ctl}$ vs F-CRF$^{HcrtR1\text{-}KO}$ p = 0.0441, *F-CRF$^{HcrtR1\text{-}Ctl}$ vs M-CRF$^{HcrtR1\text{-}Ctl}$ p = 0.0176. **B,** sEPSC frequency. Effect of Sex x Genotype p = 0.0189, *F-CRF$^{HcrtR1\text{-}Ctl}$ vs F-CRF$^{HcrtR1\text{-}KO}$ p = 0.0104, *F-CRF$^{HcrtR1\text{-}KO}$ vs M-CRF$^{HcrtR1\text{-}KO}$ p = 0.0346. **C - D,** Amplitude and frequency of spontaneous inhibitory postsynaptic currents (sIPSC) did not significantly differ across groups. **E - H,** Representative traces of sEPSC and sIPSC recordings for each sex/genotype. N = 11-16 cells/3 animals per sex per genotype.

### Supplementary Figure 5

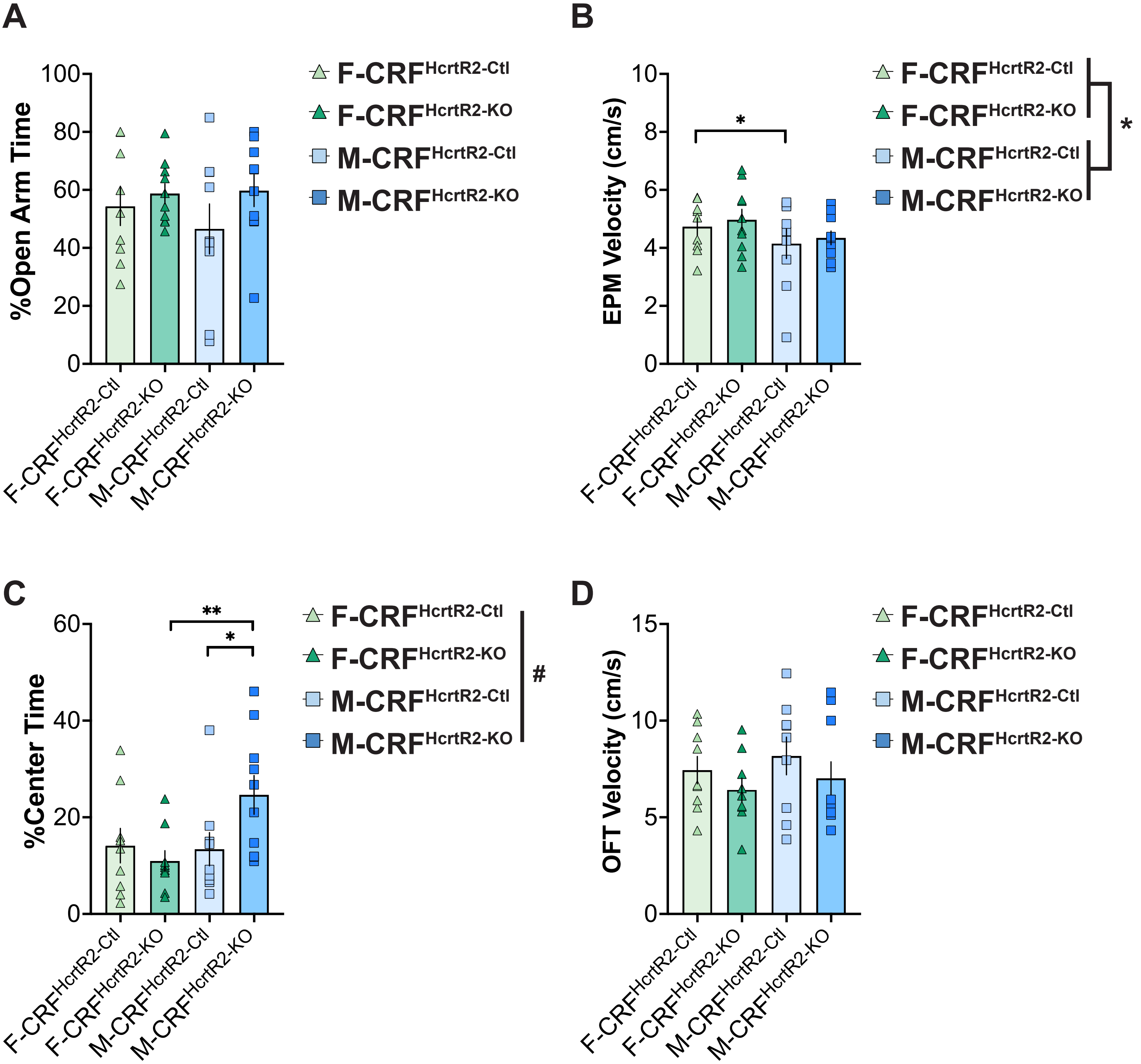

**Figure S5. HcrtR2 deletion in CRF neurons produced sex-specific impact on anxiety-like behavior in alcohol-naive mice.**

**A,** Percent time spent in the open arms in EPM tests from alcohol-naive female control (F-$CRF^{HcrtR2\text{-}Ctl}$), male control (M-$CRF^{HcrtR2\text{-}Ctl}$), female knockout (F-$CRF^{HcrtR2\text{-}KO}$), and male knockout (M-$CRF^{HcrtR2\text{-}KO}$) mice. **B,** Velocity during the EPM test across 4 groups. Effect of Sex p = 0.0225, Genotype p = 0.0595, *F-$CRF^{HcrtR2\text{-}Ctl}$ vs M-$CRF^{HcrtR2\text{-}Ctl}$ p = 0.0268. **C,** Percent time spent in the center zone in OFT from four groups of alcohol-naive mice. Effect of Sex x Genotype p = 0.0423, Sex p = 0.0665, *M-$CRF^{HcrtR2\text{-}Ctl}$ vs M-$CRF^{HcrtR2\text{-}KO}$ p = 0.0245, **F-$CRF^{HcrtR2\text{-}KO}$ vs M-$CRF^{HcrtR2\text{-}KO}$ p = 0.0071. **D,** Velocity in the OFT across 4 groups. N = 9-10 animals per sex per genotype.

### Supplementary Figure 6

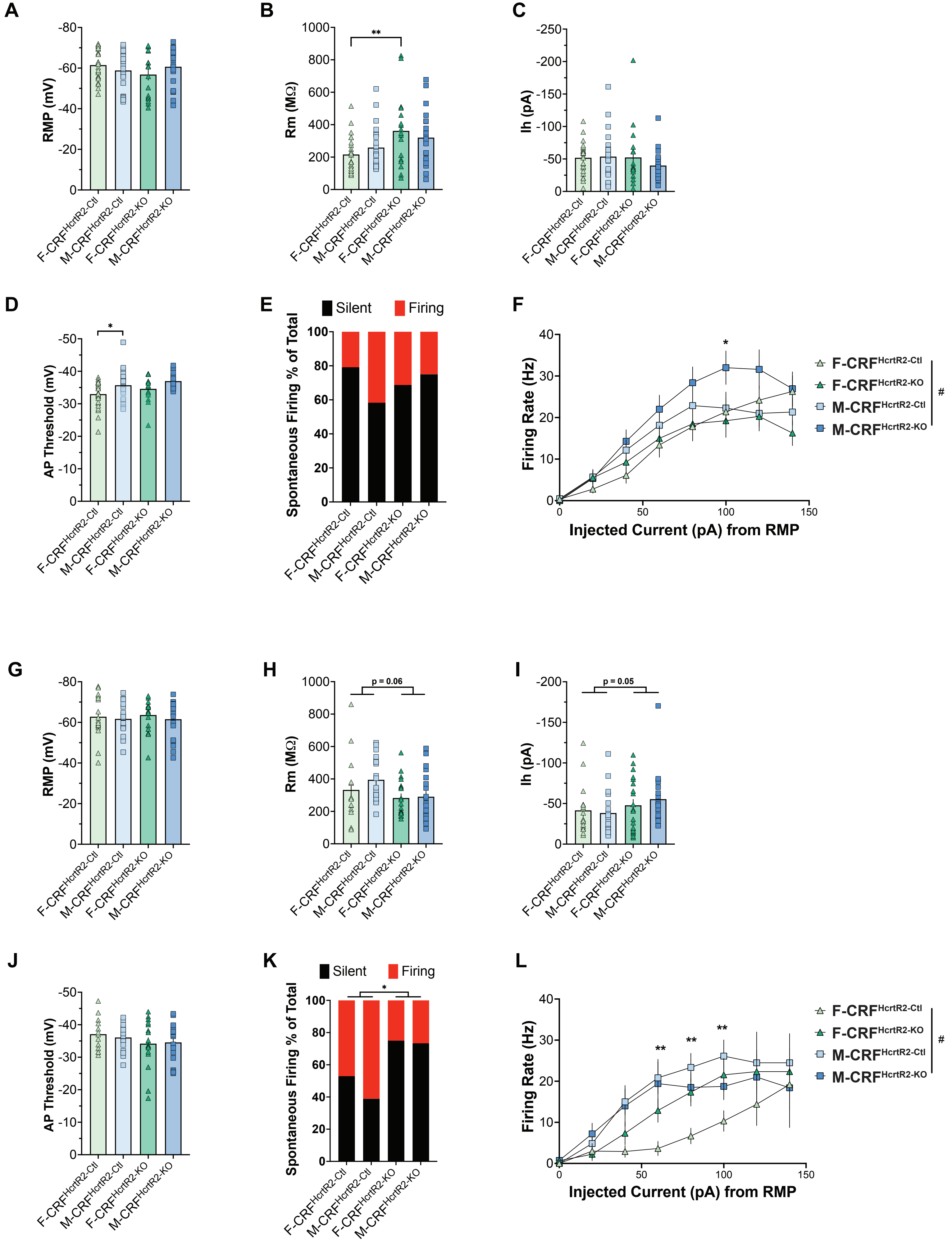

**Figure S6. HcrtR2 deletion in CRF neurons produced sex-specific effects on BNST excitability.**

**A - F,** Effect of CRF-specific HcrtR2 deletion on BNST excitability in alcohol-naive mice. **A,** RMP of BNST neurons in four groups of alcohol-naive mice. **B,** Rm of BNST neurons differed between genotypes ($p = 0.0041$), **F-$CRF^{HcrtR2-Ctl}$ vs F-$CRF^{HcrtR2-KO}$ $p = 0.0051$. **C,** Ih did not vary between groups. **D,** AP thresholds differed between sexes ($p = 0.0036$), Effect of Genotype $p = 0.0912$. *F-$CRF^{HcrtR2-Ctl}$ vs M-$CRF^{HcrtR2-Ctl}$ $p = 0.0160$, F-$CRF^{HcrtR2-KO}$ vs M-$CRF^{HcrtR2-KO}$ $p = 0.0689$. **E,** Spontaneous firing was similar across groups. **F,** CRF-specific deletion of HcrtR2 led to higher evoked firing rate in alcohol-naive males. Effect of Sex x Genotype $p = 0.0005$, Sex $p = 0.0001$, *F-$CRF^{HcrtR2-KO}$ vs M-$CRF^{HcrtR2-KO}$ $p = 0.0312$. **G - L,** Effect of CRF-specific HcrtR2 deletion on BNST excitability in alcohol-experienced mice. **G, J,** RMP and AP threshold were similar across groups. **H,** A strong trend of genotype on Rm ($p = 0.0580$). **I,** Similarly, a strong trend of genotype on Ih ($p = 0.0529$). **K,** CRF-specific HcrtR2 deletion enhanced spontaneous firing in alcohol-experience mice. Effect of Genotype $p = 0.0191$. **L,** Sex-dependent effect of CRF-specific deletion of HcrtR2 on BNST evoked firing rates. Effect of Sex x Genotype $p = 0.0021$, Sex $p < 0.0001$, F-$CRF^{HcrtR2-Ctl}$ vs M-$CRF^{HcrtR2-Ctl}$ **$p = 0.0024$, 0.0024, 0.0048.

### Supplementary Figure 7

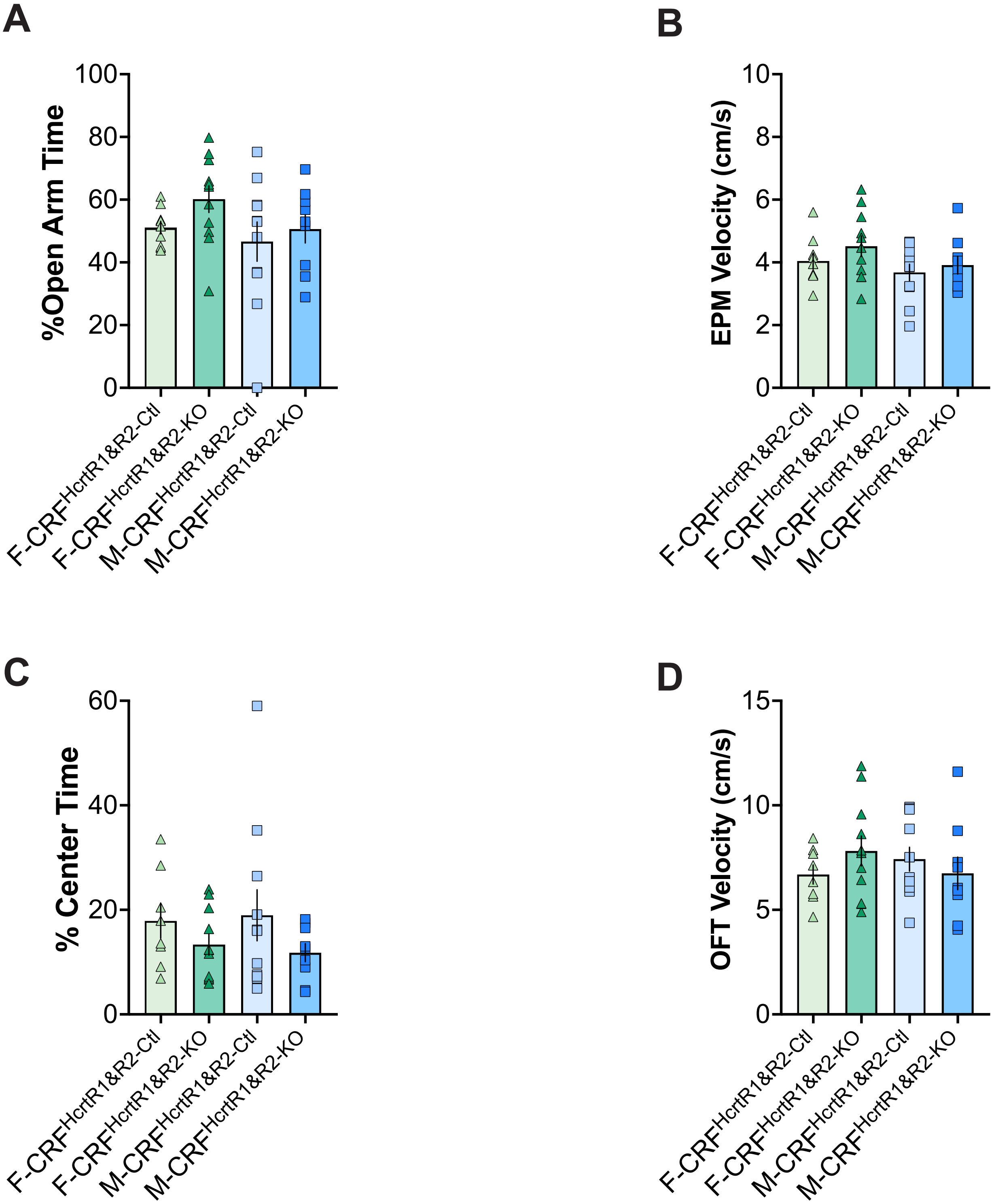

**Figure S7. Double deletion of both HcrtRs in CRF neurons did not significantly impact anxiety-like behavior in alcohol-naive mice.**

**A,** Percent time spent in the open arms in EPM tests from alcohol-naive female control (F-CRF$^{HcrtR1\&R2\text{-}Ctl}$), male control (M-CRF$^{HcrtR1\&R2\text{-}Ctl}$), female knockout (F-CRF$^{HcrtR1\&R2\text{-}KO}$), and male knockout (M-CRF$^{HcrtR1\&R2\text{-}KO}$) mice. **B,** Velocity during the EPM test across 4 groups. **C,** Percent time spent in the center zone in OFT from four groups of alcohol-naive mice. Effect of Genotype p = 0.0901. **D,** Velocity in the OFT across 4 groups. N = 9-11 animals per sex per genotype.

### Supplementary Figure 8

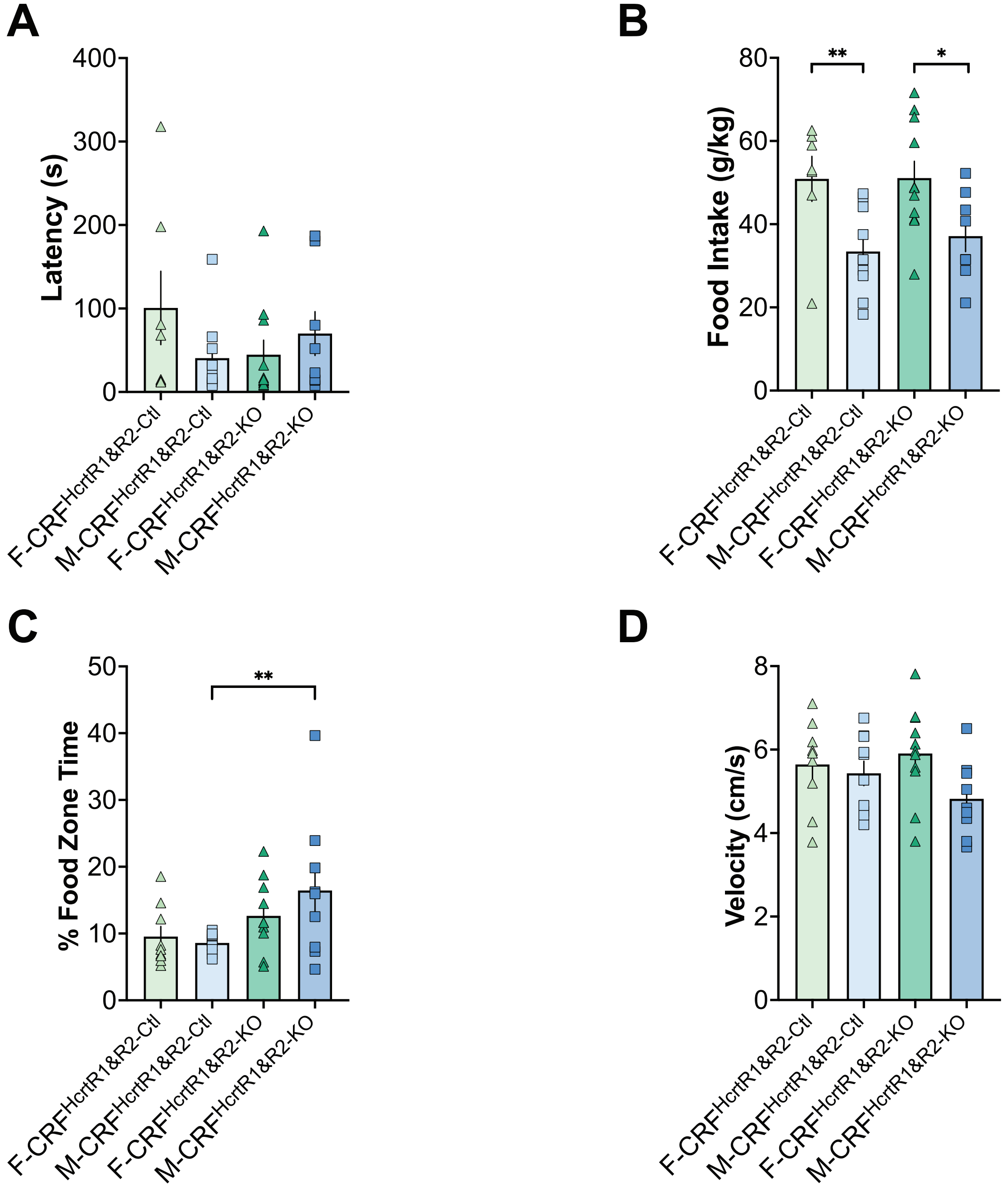

**Figure S8. Double deletion of both HcrtRs in CRF neurons produced anxiolytic effects during protracted withdrawal in the NSFT.**

**A,** Latency to investigate food in novelty-suppressed feeding test (NSFT) during protracted withdrawal after 8 weeks of 2BC IA paradigm from female control (F-$CRF^{HcrtR1\&R2\text{-}Ctl}$), male control (M-$CRF^{HcrtR1\&R2\text{-}Ctl}$), female knockout (F-$CRF^{HcrtR1\&R2\text{-}KO}$), and male knockout (M-$CRF^{HcrtR1\&R2\text{-}KO}$) mice. **B,** Food intake within 2 hours after NSFT from four groups of animals. Effect of Sex p = 0.0006, **F-$CRF^{HcrtR2\text{-}Ctl}$ vs M-$CRF^{HcrtR2\text{-}Ctl}$ p = 0.0066, *F-$CRF^{HcrtR2\text{-}KO}$ vs M-$CRF^{HcrtR2\text{-}KO}$ p = 0.0193. **C,** Percentage of time spent in the food zone during NSFT from four groups. Effect of Genotype p = 0.0098, **M-$CRF^{HcrtR1\&R2\text{-}Ctl}$ vs M-$CRF^{HcrtR1\&R2\text{-}KO}$ p = 0.0097. **D,** Velocity during NSFT across groups. Effect of Sex p = 0.0541. N = 9-11 animals per sex per genotype.
